# Native Mass Spectrometry of Complexes Formed by Molecular Glues Reveals Stoichiometric Rearrangement of E3 Ligases

**DOI:** 10.1101/2023.02.03.526954

**Authors:** Cara Jackson, Rebecca Beveridge

## Abstract

In this application of native mass spectrometry (nMS) to investigate complexes formed by molecular glues (MGs), we have demonstrated its efficiency in delineating stoichiometric rearrangements of E3 ligases that occur during targeted protein degradation (TPD). MGs stabilise interactions between an E3 ligase and a protein of interest (POI) targeted for degradation, and these ternary interactions are challenging to characterise. We have shown that nMS can unambiguously identify complexes formed between the CRBN:DDB1 E3 ligase and the POI GSPT1 upon the addition of lenalidomide, pomalidomide or thalidomide. Ternary complex formation was also identified involving the DCAF15:DDA1:DDB1 E3 ligase in the presence of MG (E7820 or indisulam) and POI RBM39. Moreover, we uncovered that the DCAF15:DDA1:DDB1 E3 ligase self-associates into dimers and trimers when analysed alone at low salt concentrations (100 mM ammonium acetate) which dissociate into single copies of the complex at higher salt concentrations (500 mM ammonium acetate), or upon the addition of MG and POI, forming a 1:1:1 ternary complex. This work demonstrates the strength of nMS in TPD research, reveals novel binding mechanisms of the DCAF15 E3 ligase, and highlights the potential effect of salt concentrations on protein complexes during structural analysis.

## Introduction

Protein-protein interactions (PPIs) play pivotal roles in many cellular processes and are therefore regarded as promising targets for drug discovery [1]. The classic approach of targeting PPIs has been to inhibit their formation, often with the use of small molecules [2, 3] or engineered peptides [4]. Additionally, the use of PPI stabilisers has also garnered significant attention [5], especially in the area of targeted protein degradation (TPD). Here, small molecules such as proteolysis-targeting chimeras (PROTACs) [6] or Molecular Glues (MGs) [7] are used to stabilise the interaction between an E3 ligase and a protein of interest (POI) that has been targeted for degradation by the cell (Figure 1). This draws the proteins into close spatial proximity, resulting in ubiquitination of the POI by the E3 ligase and subsequent proteasomal degradation of the POI. TPD is currently in an era of significant growth, and many innovative technologies are in development to address challenges in the approaches. These challenges include expanding the availability of E3 ligases that can be hijacked in TPD [8], achieving selectivity of a degrader to a specific POI [9], and the availability of analytical methodologies to directly measure the formation of ternary complexes formed between the E3 ligase, the glue, and the POI [10, 11].

**Figure 1.**
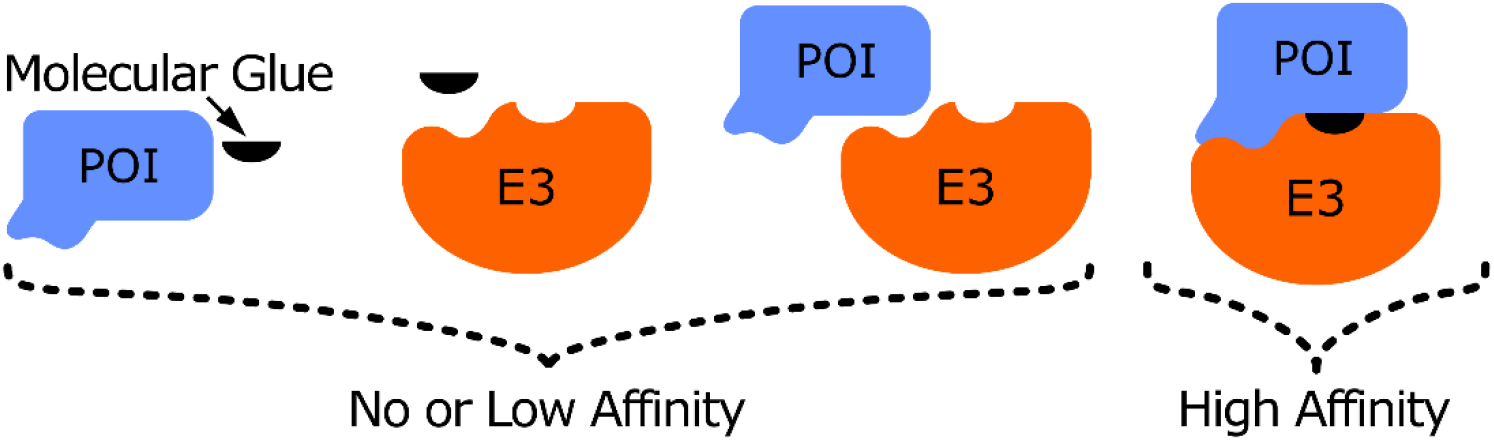
Schematic depicting the binding mechanism of molecular glues (MGs). In a typical system, interactions between the POI/MG, the E3/MG or the POI/E3 have no or low affinity, whereas the combination of all three components results in a high affinity complex.

Made infamous as the teratogenic morning sickness medication, the immunomodulatory drug (IMiD) thalidomide has gained newfound therapeutic use in the treatment of multiple myeloma [12]. Thalidomide and its derivatives lenalidomide and pomalidomide act as molecular glues between the E3 ubiquitin ligase consisting of cereblon (CRBN) and damaged DNA binding protein 1 (DDB1) (CRBN:DDB1), and various target POIs. In addition to these IMiD MGs, an additional group named the splicing inhibitor sulfonamides (SPLAMs) are becoming widely used in clinical trials, either as a single agent or in combination with other treatments [13]. E7820 and indisulam are the most documented SPLAMs, used to recruit RNA binding protein 39 (RBM39) for degradation by the E3 ubiquitin ligase consisting of DDB1 and CUL4 associated factor 15 (DCAF15), DET1 and DDB1 associated protein 1 (DDA1), and damaged DNA binding protein 1 (DDB1) (DCAF15:DDA1:DDB1). RBM39 and DCAF15 are known to have a low affinity for each other, but SPLAM molecular glues can greatly enhance the strength of this interaction [14].

Native mass spectrometry (nMS) is a means of analysing both nondenatured proteins [15, 16] and protein complexes in their native state [17, 18]. Here, large proteins can be transferred from solution into the gas phase as multiply charged ions, in a process known as electrospray ionization (ESI) [19]. nMS tends to use nanoelectrospray ionization (nESI) [20], which produces finer droplets, allowing for native topologies, stoichiometries, and non-covalent interactions to remain while transferring proteins from a nondenaturing buffer solution, commonly ammonium acetate (AmAc) [21], into the gas phase [22].

Herein, nMS is demonstrated as a label-free, sensitive, and straightforward tool for detecting the stoichiometry of intact protein complexes formed with MGs. nMS is effective in separating individual species that exist in a stoichiometric mixture, capturing transient interactions, and comparing stability of complexes [17, 23, 24]. Whilst nMS has previously been used to predict the efficacy of PROTACs [25] and to analyse complexes formed between MGs and model peptides [26], this is the first example of its application to MGs and multimeric E3 complexes. We have demonstrated its efficacy using disease relevant drugs and targets; two E3 ligase proteins (CRBN:DDB1 and DCAF15:DDA1:DDB1), five molecular glues (lenalidomide, thalidomide, pomalidomide, E7820, and indisulam, Table S1), and two POIs (GSPT1 and RBM39).

## Results

### nMS identifies IMiD-mediated interactions between CRBN:DDB1 and GSPT1

We first sought to investigate the complexes formed by the IMiD MGs lenalidomide, thalidomide and pomalidomide with the CRBN:DDB1 E3 ligase and the GSPT1 POI. CRBN:DDB1 and GSPT1 (5μM each) were sprayed in a mixture from 100 mM AmAc in the absence and presence of the MG lenalidomide (Figure 2A and B, respectively). AmAc is the most popular solvent in nMS as it is volatile and provides the required pH (6.8) for native protein analysis [21]. In the absence of MGs, no interactions are observed between GSPT1 and CRBN:DDB1 (Figure 2A). Here, CRBN:DDB1 presents in charge states 17+ to 23+, monomeric DDB1 presents in charge states 14+ to 19+ and the POI GSPT1 presents in three charge states, from 8+ to 10+. Upon the addition of lenalidomide at 100 μM (Figure 2B), new peaks corresponding to the E3:MG:POI complex can be observed in six charge states from 20+ to 25+. The same is observed upon the addition of additional MGs thalidomide and pomalidomide (Figure S1).

**Figure 2.**
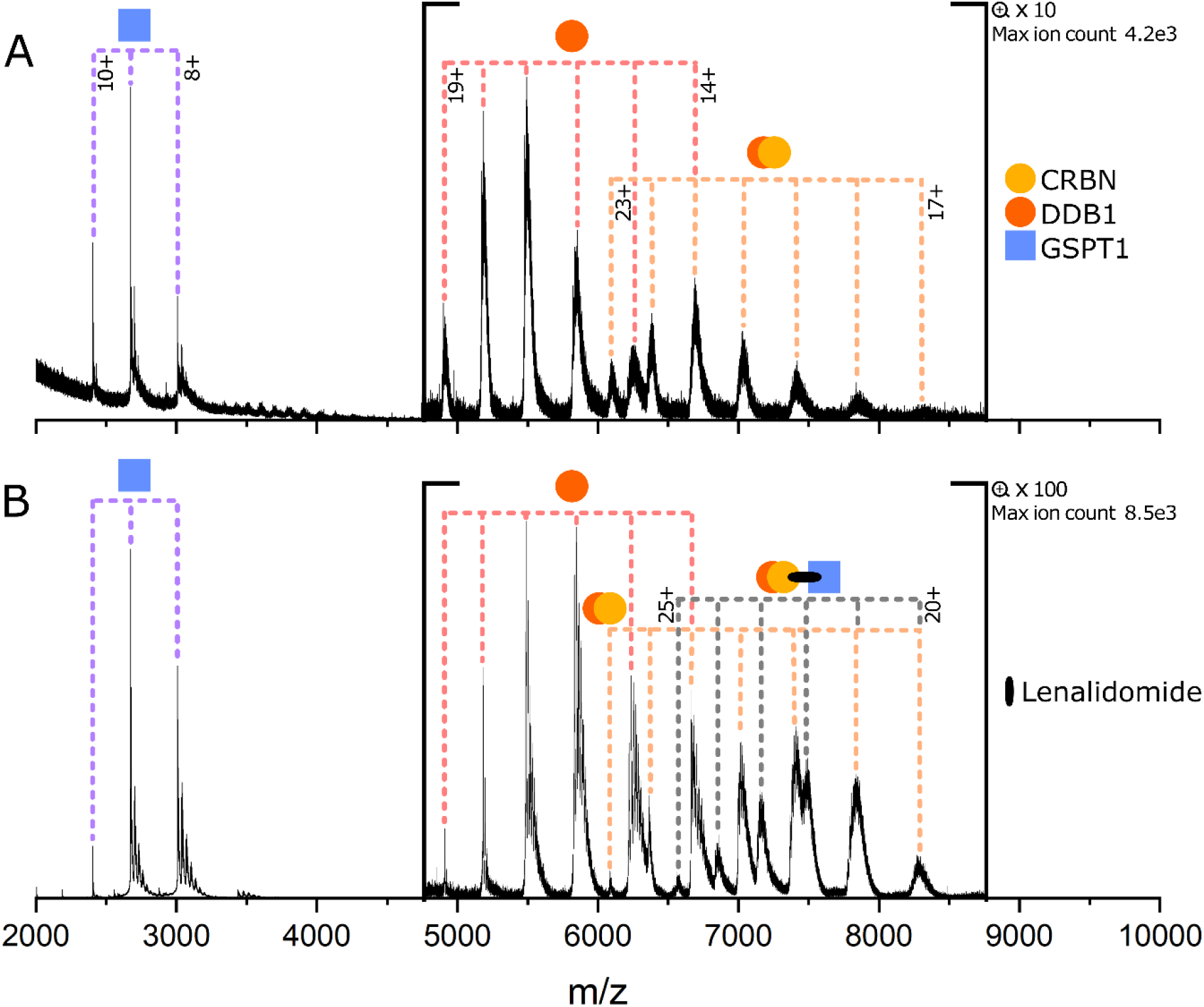
(A) GSPT1 + CRBN:DDB1 (B) GSPT1 + CRBN:DDB1 + lenalidomide. Protein concentrations are 5 μM and lenalidomide is 100 μM when present.

Lenalidomide was introduced to the E3:POI mixture at lower concentrations of 50 μM and 5 μM, and ternary complex is still observed in both cases, albeit at a low relative intensity (Figure S2). No peaks corresponding to binary species (E3:MG or POI:MG) were observed in the mixture of all three components (Figure 2, S1, S2), nor when CRBN:DDB1 + MG was sprayed in the absence of GSPT1 (Figure S3). IMiDs have previously been shown to bind CRBN with weak KDs of 10-65 μM, which we would not expect to observe with nMS [27]. The measured mass of all proteins used in this study is given in Table S2.

### The DCAF15 complex forms dimers and trimers in the absence of MG and POI

We next turned our attention to the DCAF15-targeting SPLAM MG E7820 (Figure 3). The DCAF15:DDA1:DDB1 complex was used, lacking the proline-rich, atrophin-homology domain of DCAF15 (amino acids 276–383) which has been used in previous studies, and we refer to here as the DCAF15 complex [14]. The POI used is the RRM2 domain of RBM39, which we refer to as RBM39 hereafter. In this case, the control experiment containing RBM39 (4+ to 6+) and the DCAF15 complex in the absence of MG yielded extremely surprising results. Monomeric DCAF15 complex (charge states 19+ to 24+) is only observed to a very low extent, and most of the signal intensity of this species corresponds to dimers and trimers of the DCAF15 complex (Figure 3A). The dimer of the DCAF15 complex presents in nine charge states from 29+ to 37+ and the trimer also presents in nine charge states, 36+ to 44+. Unbound DDB1 is also present in charge states 14+ to 18+, and the DDA1:DDB1 dimer is present in charge states 15+ to 17+.

**Figure 3.**
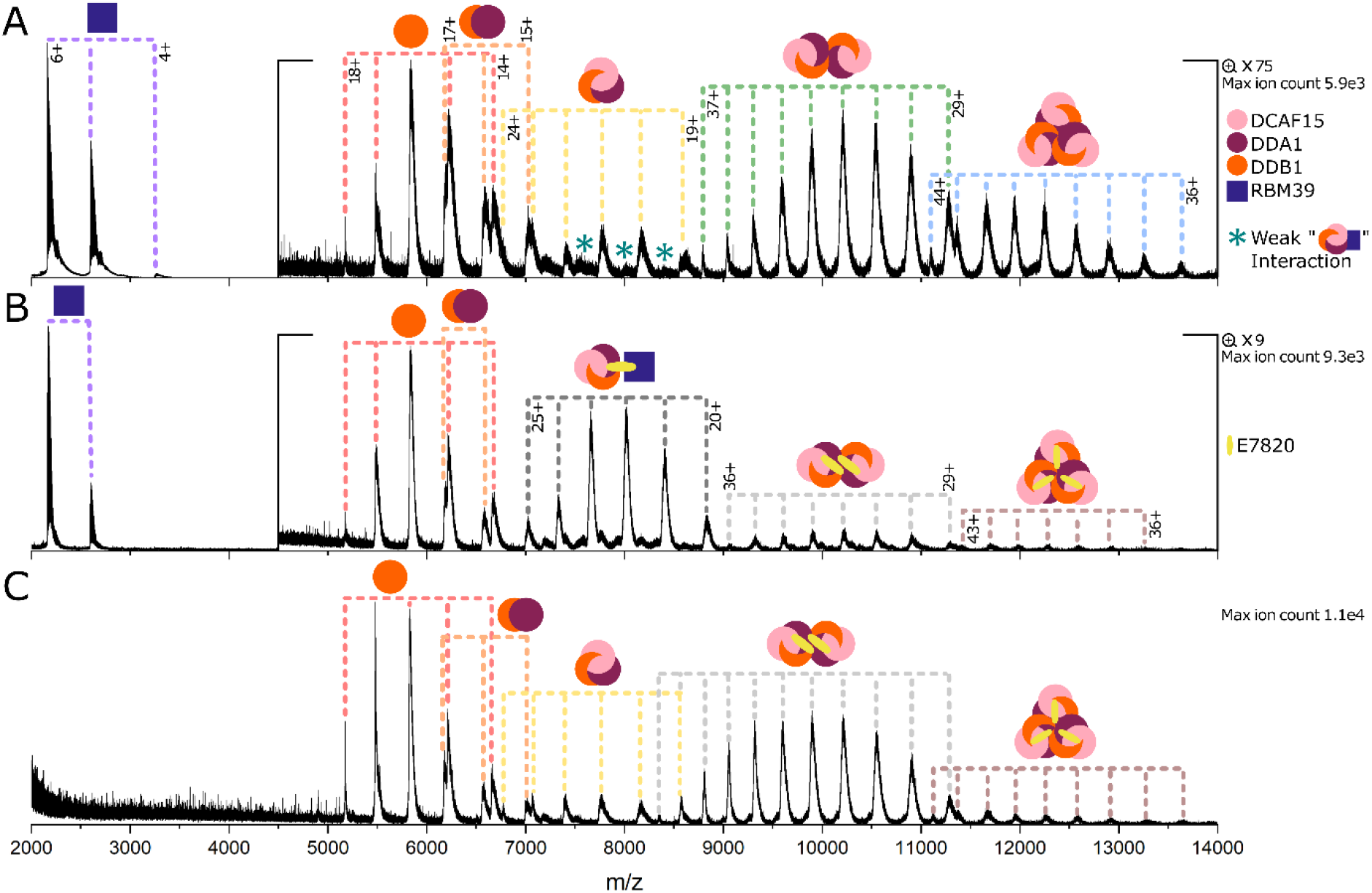
(A) RBM39 + DCAF15 complex (DCF15:DDA1:DDB1). (B) RBM39 + DCAF15 complex + E7820. (C) DCAF15 complex + E7820. Protein concentrations are 5 μM and E7820 is 100 μM when present.

To identify whether this multimerization is concentration dependent, the DCAF15 complex was analysed alone at concentrations of 5 μM and 2.5 μM (Figure S4) which yielded very similar complex distributions. This indicated that formation of the dimers and trimers is not concentration dependent. Upon close observation, peaks can also be identified corresponding to the 1:1 complex between the DCAF15 complex and RBM39, as indicated by asterisks in Figures 3, S5 and S6. This is in agreement with the literature, as RBM39 is known to be a native substrate of the DCAF15 complex, and a weak interaction of 4-6 μM has previously been measured between the two species [14].

Upon addition of the MG E7820 (Figure 3B) to the DCAF15 complex and the RBM39, a stoichiometric rearrangement of the DCAF15 complex occurs and the main signal intensity now corresponds to a single copy of the complex bound to E7820 and RBM39 in a 1:1:1 stoichiometry (20+ to 25+). Low amounts of dimeric DCAF15 complex remain present bound to two molecules of E7820 (Figure 3B), and the trimeric complex is almost completely eradicated. To investigate whether this stoichiometric rearrangement of the DCAF15 complex is due to the MG or the POI, the DCAF15 complex and E7820 were sprayed together in the absence of RBM39. In this case, the DCAF15 complex is mainly observed as dimers bound to E7820 in a 2:2 stoichiometry (Figure 3C and S7). Very low signal for the trimer of the DCAF15 complex is observed bound to three E7820 molecules (3:3 complex), but the intensity of the trimer is much lower than for the DCAF15 complex in the absence of MG (Figures 3A and S4). We therefore hypothesise that the MG destabilises the DCAF15 complex trimer and causes preference for the dimer. No interaction was seen between the monomeric DCAF15 complex and E7820, which has been previously measured to have a K_D_ > 50 μM [28]. This is a weak interaction that would not be expected to be observed with nMS, but the fact that the 2:2 complex is seen suggests that the MGs bind at the interface of the dimer and have a stronger interaction than with the monomer (Figure S7). Equivalent data for the indisulam MG is shown in Figure S5.

As a complementary approach to nMS and to consolidate these findings, size exclusion chromatography (SEC) was employed to compare the interactions of the DCAF15 complex, RBM39 and E7820 (Figure S8). This confirmed the presence of higher order oligomers of the DCAF15 complex when analysed alone, as well as the formation of the ternary complex in the presence of E7820 and RBM39.

### Interactions of the DCAF15 complex are strongly regulated by salt concentration

To further investigate this unexpected oligomerisation of the DCAF15 complex, it was analysed from solutions of increased AmAc concentration to examine the effect of ionic strength on the interactions (Figure 4). When ionised from a solution of 100 mM AmAc (Figure 4a), the DCAF15 complex is present in a mixture of monomers, dimers and trimers, with the signal corresponding to the dimer being the most intense. Upon increasing the AmAc concentration of the starting solution to 500 mM (Figure 4B), the majority of the signal corresponds to monomeric DCAF15, with just a small amount of signal corresponding to the dimeric form and no signal remaining for the trimer. This means that higher AmAc concentrations, and therefore higher ionic strength, shifts the DCAF15 complex from dimers and trimers towards the monomeric form. To determine whether the DCAF15 is still able to form a ternary complex in these conditions it was analysed in the presence of E7820 and RBM39 (Figure 4C) which resulted in the formation of ternary complex. However, in these high salt conditions, there remains a portion of unbound DCAF15 complex, which is not present when analysed from the low salt solution (Figure 3B). This implies that the higher ionic strength solution is disrupting interactions between the DCAF15 complexes, as well as with the RBM39 POI.

**Figure 4.**
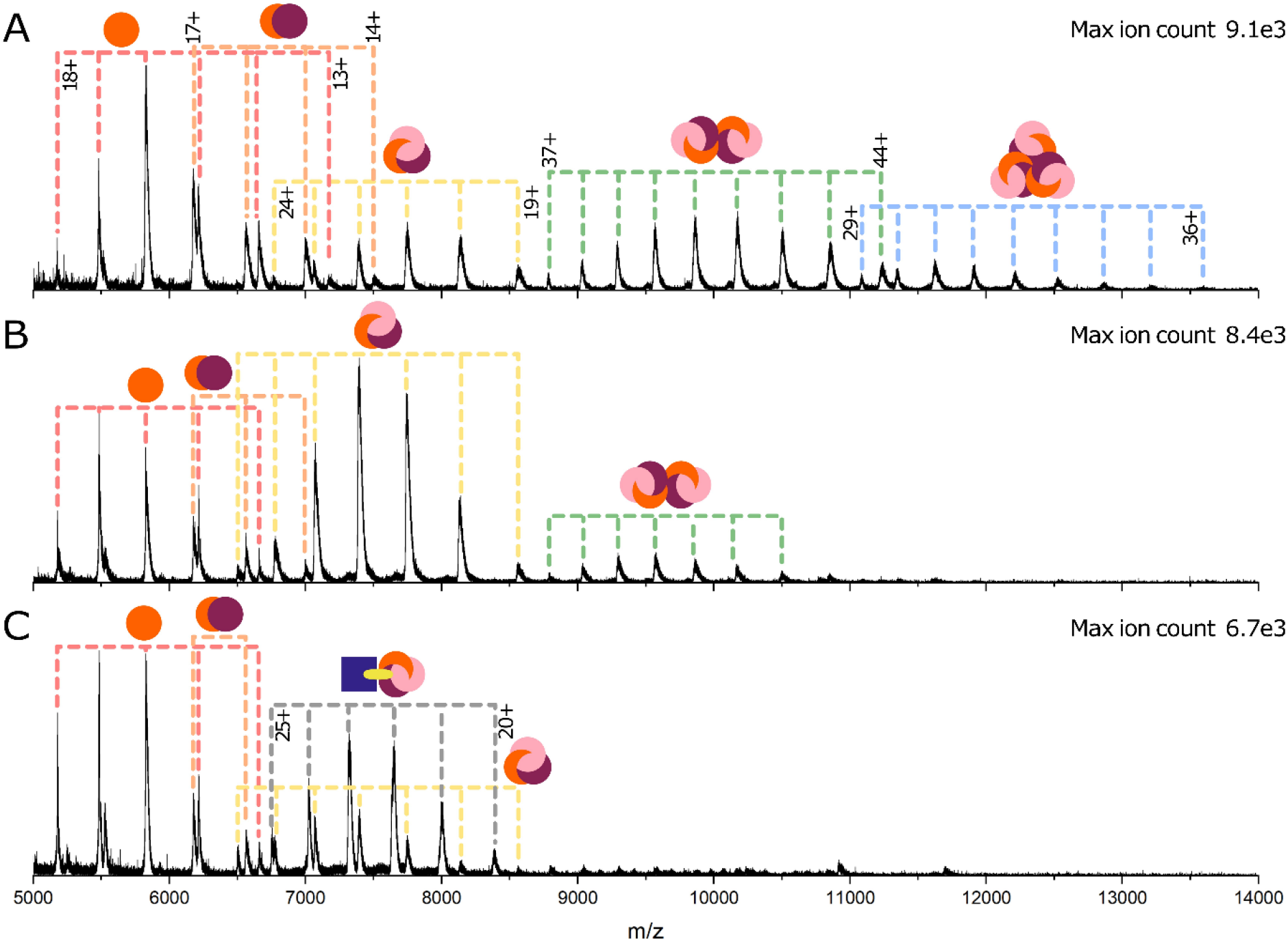
(A) DCAF15 complex ionised from 100 mM AmAc. (B) DCAF15 complex ionised from 500 mM AmAc. (C) DCAF15 complex + RBM39 + E7820 complex ionised from 500 mM AmAc. Protein concentrations are 5 μM and E7820 is 100 μM when present.

This dissociation of DCAF15 multimers in response to high salt concentration is not a feature displayed across all proteins. In a study by Gavriilidou et al. [29], it was shown that increasing AmAc concentrations up to 500 mM promoted tetramer formation of Concanavalin A over the dimeric form. In this same study it was shown that protein-ligand interactions are affected differently by high AmAc concentration, with interactions increasing in affinity between Lysozyme-NaG3 (Tri-N-acetylchitotriose) and Trypsin–Pefabloc, and interactions decreasing in affinity between Carbonic Anhydrase II-chlorothiazide and β-Lactoglobulin-Lauric Acid. Despite the importance of AmAc solutions in native mass spectrometry to preserve protein structure there remains little evidence in the literature regarding its optimum concentration. However, this significant change in the ability of the DCAF15 complex to multimerise and form ternary complexes may be a feature of the protein that contributes to its activity and regulation, and this hypothesis will be addressed in subsequent research.

## Conclusions

In summary, nMS has been demonstrated as an asset for the determination of MG ternary complex formation, successfully and clearly showing the presence of two ternary complexes between disease relevant proteins. It is a sensitive, fast, label-free technique requiring low sample consumption. This work has shown that nMS can show these complexes with intact proteins, in addition to the model peptides which have previously been used [26]. nMS was able to directly show the existence of the IMiD ternary complexes, which will be beneficial in the screening of small molecule libraries for further glues that modulate this interaction.

An unexpected and important outcome of this study was that the DCAF15 complex self-associates into dimers and trimers, which are disrupted either in the presence of (i) E7820/indisulam and RBM39 or (ii) higher salt concentrations. Despite this E3 ligase being extensively studied in structural biology due to its potential role in TPD, oligomerisation has not previously been reported, to our knowledge. Such findings are paramount in understanding E3 ligases for their manipulation in TPD, and we expect that analysis of complexes via nMS will eventually be routine in guiding drug design.

## Acknowledgements

This work was funded by Triana Biomedicines Inc, Waltham, MA, USA. RB acknowledges support of a UKRI Future Leaders Fellowship (Grant Reference MR/T020970/1) and the University of Strathclyde for a Chancellor’s Fellowship (2020-2022). CJ is supported by an EPSRC studentship. AstraZeneca is thanked for providing CJ with a CASE top-up. The authors acknowledge the MVLS Structural Biology and Biophysical Characterisation Facility, University of Glasgow, for the SEC analysis.

## References

1. Scott, D.E., A.R. Bayly, C. Abell, and J. Skidmore, Small molecules, big targets: drug discovery faces the protein–protein interaction challenge. Nature Reviews Drug Discovery, 2016. 15(8): p. 533–550.

2. Gorczynski, M.J., J. Grembecka, Y. Zhou, Y. Kong, L. Roudaia, M.G. Douvas, M. Newman, I. Bielnicka, G. Baber, T. Corpora, J. Shi, M. Sridharan, R. Lilien, B.R. Donald, N.A. Speck, M.L. Brown, and J.H. Bushweller, Allosteric Inhibition of the Protein-Protein Interaction between the Leukemia-Associated Proteins Runx1 and CBFβ. Chemistry & Biology, 2007. 14(10): p. 1186–1197.

3. Yin, X., C. Giap, J.S. Lazo, and E.V. Prochownik, Low molecular weight inhibitors of Myc–Max interaction and function. Oncogene, 2003. 22(40): p. 6151–6159.

4. Dietrich, L., B. Rathmer, K. Ewan, T. Bange, S. Heinrichs, T.C. Dale, D. Schade, and T.N. Grossmann, Cell Permeable Stapled Peptide Inhibitor of Wnt Signaling that Targets β-Catenin Protein-Protein Interactions. Cell Chemical Biology, 2017. 24(8): p. 958–968.e5.

5. Thiel, P., M. Kaiser, and C. Ottmann, Small-Molecule Stabilization of Protein–Protein Interactions: An Underestimated Concept in Drug Discovery? Angewandte Chemie International Edition, 2012. 51(9): p. 2012–2018.

6. Sakamoto, K.M., K.B. Kim, A. Kumagai, F. Mercurio, C.M. Crews, and R.J. Deshaies, Protacs: Chimeric molecules that target proteins to the Skp1–Cullin–F box complex for ubiquitination and degradation. Proceedings of the National Academy of Sciences, 2001. 98(15): p. 8554–8559.

7. Słabicki, M., Z. Kozicka, G. Petzold, Y.-D. Li, M. Manojkumar, R.D. Bunker, K.A. Donovan, Q.L. Sievers, J. Koeppel, D. Suchyta, A.S. Sperling, E.C. Fink, J.A. Gasser, L.R. Wang, S.M. Corsello, R.S. Sellar, M. Jan, D. Gillingham, C. Scholl, S. Fröhling, T.R. Golub, E.S. Fischer, N.H. Thomä, and B.L. Ebert, The CDK inhibitor CR8 acts as a molecular glue degrader that depletes cyclin K. Nature, 2020. 585(7824): p. 293–297.

8. Kramer, L.T. and X. Zhang, Expanding the landscape of E3 ligases for targeted protein degradation. Current Research in Chemical Biology, 2022. 2: p. 100020.

9. Smith, B.E., S.L. Wang, S. Jaime-Figueroa, A. Harbin, J. Wang, B.D. Hamman, and C.M. Crews, Differential PROTAC substrate specificity dictated by orientation of recruited E3 ligase. Nature Communications, 2019. 10(1): p. 131.

10. Gadd, M.S., A. Testa, X. Lucas, K.-H. Chan, W. Chen, D.J. Lamont, M. Zengerle, and A. Ciulli, Structural basis of PROTAC cooperative recognition for selective protein degradation. Nature Chemical Biology, 2017. 13(5): p. 514–521.

11. Roy, M.J., S. Winkler, S.J. Hughes, C. Whitworth, M. Galant, W. Farnaby, K. Rumpel, and A. Ciulli, SPR-Measured Dissociation Kinetics of PROTAC Ternary Complexes Influence Target Degradation Rate. ACS Chemical Biology, 2019. 14(3): p. 361–368.

12. Yamamoto, J., T. Ito, Y. Yamaguchi, and H. Handa, Discovery of CRBN as a target of thalidomide: a breakthrough for progress in the development of protein degraders. Chemical Society Reviews, 2022. 51(15): p. 6234–6250.

13. Milojkovic Kerklaan, B., S. Slater, M. Flynn, A. Greystoke, P.O. Witteveen, M. Megui-Roelvink, F. de Vos, E. Dean, L. Reyderman, L. Ottesen, M. Ranson, M.P.J. Lolkema, R. Plummer, R. Kristeleit, T.R.J. Evans, and J.H.M. Schellens, A phase I, dose escalation, pharmacodynamic, pharmacokinetic, and food-effect study of α2 integrin inhibitor E7820 in patients with advanced solid tumors. Investigational New Drugs, 2016. 34(3): p. 329–337.

14. Du, X., O.A. Volkov, R.M. Czerwinski, H. Tan, C. Huerta, E.R. Morton, J.P. Rizzi, P.M. Wehn, R. Xu, D. Nijhawan, and E.M. Wallace, Structural Basis and Kinetic Pathway of RBM39 Recruitment to DCAF15 by a Sulfonamide Molecular Glue E7820. Structure, 2019. 27(11): p. 1625–1633.e3.

15. Karas, M., U. Bahr, and T. Dülcks, Nano-electrospray ionization mass spectrometry: addressing analytical problems beyond routine. Fresenius’ Journal of Analytical Chemistry, 2000. 366(6-7): p. 669–676.

16. Leney, A.C. and A.J.R. Heck, Native Mass Spectrometry: What is in the Name? Journal of the American Society for Mass Spectrometry, 2017. 28(1): p. 5–13.

17. Beveridge, R., L.G. Migas, K.A.P. Payne, N.S. Scrutton, D. Leys, and P.E. Barran, Mass spectrometry locates local and allosteric conformational changes that occur on cofactor binding. Nature Communications, 2016. 7(1): p. 12163.

18. Bellamy-Carter, J., M. Mohata, M. Falcicchi, J. Basran, Y. Higuchi D.R. G., and A.C. Leney, Discovering protein–protein interaction stabilisersby native mass spectrometry. Chemical Science, 2021. 12: p. 10724–10731.

19. Fenn, J.B., M. Mann, C.K. Meng, S.F. Wong, and C.M. Whitehouse, Electrospray Ionization for Mass Spectrometry of Large Biomolecules. Science., 1989. 246(4926): p. 64–71.

20. Wilm, M. and M. Mann, Analytical Properties of the Nanoelectrospray Ion Source. Analytical Chemistry, 1996. 68(1): p. 1–8.

21. Konermann, L., Addressing a Common Misconception: Ammonium Acetate as Neutral pH “Buffer” for Native Electrospray Mass Spectrometry. Journal of the American Society for Mass Spectrometry, 2017. 28(9): p. 1827–1835.

22. Sharon, M. and C.V. Robinson, The role of mass spectrometry in structure elucidation of dynamic protein complexes. Annual review of biochemistry, 2007. 76(1): p. 167–193.

23. Sugiyama, M., H. Yagi, K. Ishii, L. Porcar, A. Martel, K. Oyama, M. Noda, Y. Yunoki, R. Murakami, R. Inoue, N. Sato, Y. Oba, K. Terauchi, S. Uchiyama, and K. Kato, Structural characterization of the circadian clock protein complex composed of KaiB and KaiC by inverse contrast-matching small-angle neutron scattering. Scientific Reports, 2016. 6(1): p. 35567.

24. Cveticanin, J., T. Mondal, E.M. Meiering, M. Sharon, and A. Horovitz, Insight into the Autosomal-Dominant Inheritance Pattern of SOD1-Associated ALS from Native Mass Spectrometry. Journal of Molecular Biology, 2020. 432(23): p. 5995–6002.

25. Beveridge, R., D. Kessler, K. Rumpel, P. Ettmayer, A. Meinhart, and T. Clausen, Native Mass Spectrometry Can Effectively Predict PROTAC Efficacy. ACS Central Science, 2020. 6(7): p. 1223–1230.

26. Bellamy-Carter, J., M. Mohata, M. Falcicchio, J. Basran, Y. Higuchi, R.G. Doveston, and A.C. Leney, Discovering protein–protein interaction stabilisers by native mass spectrometry. Chemical Science, 2021. 12(32): p. 10724–10731.

27. Akuffo, A.A., A.Y. Alontaga, R. Metcalf, M.S. Beatty, A. Becker, J.M. McDaniel, R.S. Hesterberg, W.E. Goodheart, S. Gunawan, M. Ayaz, Y. Yang, M.R. Karim, M.E. Orobello, K. Daniel, W. Guida, J.A. Yoder, A.M. Rajadhyaksha, E. Schönbrunn, H.R. Lawrence, N.J. Lawrence, and P.K. Epling-Burnette, Ligand-mediated protein degradation reveals functional conservation among sequence variants of the CUL4-type E3 ligase substrate receptor cereblon. J Biol Chem, 2018. 293(16): p. 6187–6200.

28. Bussiere, D.E., L. Xie, H. Srinivas, W. Shu, A. Burke, C. Be, J. Zhao, A. Godbole, D. King, R.G. Karki, V. Hornak, F. Xu, J. Cobb, N. Carte, A.O. Frank, A. Frommlet, P. Graff, M. Knapp, A. Fazal, B. Okram, S. Jiang, P.-Y. Michellys, R. Beckwith, H. Voshol, C. Wiesmann, J.M. Solomon, and J. Paulk, Structural basis of indisulam-mediated RBM39 recruitment to DCAF15 E3 ligase complex. Nature Chemical Biology, 2020. 16(1): p. 15–23.

29. Gavriilidou, A.F.M., B. Gülbakan, and R. Zenobi, Influence of Ammonium Acetate Concentration on Receptor–Ligand Binding Affinities Measured by Native Nano ESI-MS: A Systematic Study. Analytical Chemistry, 2015. 87(20): p. 10378–10384.

