## Supplementary material for "Native Mass Spectrometry of Complexes Formed by Molecular Glues Reveals Stoichiometric Rearrangement of E3 Ligases": Methods and supplementary figures

Cara Jackson, Rebecca Beveridge
Department of Pure and Applied Chemistry, University of Strathclyde, Glasgow, UK, G1 1XL

### Materials and Methods

*Protein Expression and Purification*

E3 ligase and target proteins

Cereblon (amino acids 40-442) with an N-terminal 6xHis-TEV tag and DDB1 (amino acids 1-1140 with replacement of 396-705 by GNGNSG) were co-expressed in SF9 insect cell. After lysis in 50 mM HEPES/NaOH, 500 mM NaCl, 1 mM TCEP, Complete protease inhibitor 1 tab / 100mL, 10 mM imidazole, pH 8.0 and centrifugation to clarify the lysate, the cereblon/DDB1 complex (hereafter CRBN/DDB1) were captured on HisPur Ni-NTA resin and eluted with a step-wise gradient with imidazole. The eluted sample was treated with TEV protease to remove the His-tag and dialyzed overnight to remove imidazole. The dialyzed sample was then passed back on the Ni-NTA resin to re-capture uncleaved samples and separated His-tag while cleaved CRBN/DDB1 passed through the resin unbound. CRBN/DDB1 was then further purified using a HiTrap Q HP ion-exchange column with a salt-gradient and size-exclusion chromatography using HiLoad Superdex 26/600 in 10 mM Hepes, 240 mM NaCl, 1 mM TCEP, pH 7.0.

GSPT1 (amino acids 300-496) containing N-terminal 6xHis-MBP-TEV and C-terminal Avi tags was co-expressed with BirA in *E. coli* BL21-CodonPlus (DE3)-RIPL. The soluble fraction generated by cell lysis in 50 mM HEPES / NaOH, 300 mM NaCl, cOmplete protease inhibitor EDTA free 1 tab / 100 mL, 1 mM TCEP, 10% glycerol, pH 7.5 and centrifugation, was passed through HisPur Ni-NTA resin to capture the target protein. Proteins were eluted from the Ni-NTA resin using an imidazole gradient and were subjected to TEV protease cleavage while dialyzing overnight to remove imidazole. The dialyzed sample was then re-applied on the Ni-NTA resin to separate tag-cleaved GSPT1 from other species. GSPT1 was further purified by heparin chromatography, and by size-exclusion chromatography in 50 mM HEPES / NaOH,150 mM NaCl,1 mM TCEP, pH 7.5.

DCAF15 (amino acids 1-600 with an internal deletion of 276-383) with N-terminal 6xHis-TEV, DDB1 (amino acids 1-1140 with replacement of 396-705 by GNGNSG), and DDA1 (amino acids 1-102) were co-expressed in SF9 cells. Cells were lysed in 25 mM Tris-HCl, 300 mM NaCl, 10% Glycerol, 1 mM TCEP, 20 mM imidazole, cOmplete protease inhibitor EDTA free 1 tab / 100 mL, 0.1 mg Benzonase/1L culture, pH 7.5 by a high-pressure homogenizer (500 bar, 3 passes) and the soluble fraction was separated by high-speed centrifugation. A HisTrap HP column was used as the first step to purify the DCAF15/DDB1/DDA1 complex from the soluble lysate. Samples eluted from the first column using an imidazole gradient were then further purified by ion-exchange chromatography, followed by size-exclusion chromatography in 25 mM Tris/HCl, 300 mM NaCl, 1 mM TCEP, pH 7.5.

RBM39 (amino acids 235-331) containing N-terminal 6xHis-TEV-Avi was expressed in *E. coli* BL21 (CP). Cell pellets were re-suspended in the lysis buffer (25 mM HEPES pH 7.5, 300 mM NaCl, 1 mM TCEP) supplemented with 0.5 mM PMSF and sonicated to generate cell lysates which was then clarified by high-speed centrifugation. The supernatant was applied on a Ni-NTA Superflow resin and the bound proteins were eluted with a step-wise imidazole gradient. The eluted proteins were treated with TEV protease to release the His-Tag and was dialyzed overnight to remove imidazole. The digested and dialyzed sample treated with BirA in the presence of biotin, magnesium chloride, and ATP overnight at 4 °C to produce biotinylated RBM39. The sample was then reloaded on a Ni-NTA column to capture uncleaved proteins and liberated His tag, leaving biotinylated and tag-cleaved RBM39 in the flow-through. This fraction was collected, concentrated, and purified by size-exclusion chromatography in 25 mM HEPES pH7.5, 300 mM NaCl, 1 mM TCEP.

*Mass Spectrometry Sample Preparation*

All proteins were dialysed into a buffer solution of 200 mM ammonium acetate (Fisher Scientific, Loughborough, Leicestershire, UK) at pH 6.8. Proteins were diluted to 20 µM in 200 mM ammonium acetate. 20 µM E3 and 20 µM POI were combined 1:1 to give a mixture consisting of 10 µM E3, 10 µM POI, and 200 mM ammonium acetate. 20 mM Molecular Glue (WuXi AppTec Co., Ltd., Shanghai, China) in 100% DMSO (Sigma-Aldrich, St. Louis, MO, USA) was diluted to 200 µM in 1% DMSO using deionised water. 200 µM Molecular Glue in 1% DMSO was combined 1:1 with the 10 µM E3 + 10 µM POI in 200 mM ammonium acetate mixture to give a final analytical concentration of 5µM E3, 5 µM POI, 100 µM molecular glue, 100 mM ammonium acetate, 0.5% DMSO. Samples were diluted to analytical concentration the day of analysis.

*Mass Spectrometry*

Mass spectrometry experiments were carried out using a Waters Synapt G2-Si mass spectrometer (Waters Corporation, Manchester, UK). A nano electrospray ionisation source (nESI) was used for ionisation. Nano electrospray tips for nESI were pulled in house from thin-walled borosilicate glass capillaries (i.d. 0.78 mm, o.d. 1.0 mm) (Sutter Instrument Co., Novato, CA, USA) using a Flaming/Brown micropipette puller (Sutter Instrument Co., Novato, CA, USA). A positive potential of 0.9-1.5 Kv was applied to the solution using a thin platinum wire (d. 0.125 mm) (Goodfellow, Huntingdon, UK). Other non-default instrument settings include: sampling cone voltage 60-200 V, source offset voltage 80-150 V, collision voltage 2-4 V, trap gas flow 4-5 ml/min, source temperature 40°C.

*Size Exclusion Chromatography*

Size exclusion chromatography experiments were carried out using a Waters Alliance HPLC system (Waters Corporation, Manchester, UK) equipped with a Waters 176003596 XBridge BEH200 SEC 3.5µm 7.8x300 column (Waters Corporation, Manchester, UK). Samples were ran using a 100 mM ammonium acetate buffer solution containing 0.5% DMSO at a flow rate of 1 mL/min. Other parameters include: laser wavelength 660nm, laser power 80%, temperature 22°C, fit type Debye, fit order 1$^{st}$, calibration method BSA 1 mg/ml. The column was equilibrated with the corresponding buffer prior to the experiment. Concentrations in Figure 4 (A), (C), and (D) are DCAF15 complex (1 µM), RBM39 (1 µM), and E7820 (20 µM) when present, and in Figure 4 (B), RBM39 (5 µM). All samples were in a 100 mM ammonium acetate buffer solution containing 0.5% DMSO (concentrations were altered due to column loading constraints).

*Data Processing*

Mass spectrometry data was processed using MassLynx (Version 4.2, Waters Corporation, Manchester, UK) and mass spectra figures were produced with OriginPro (Version 2022, OriginLab Corporation, Northampton, MA, USA) and Inkscape (Version 1.2, Inkscape.org).

### Supplementary Figures

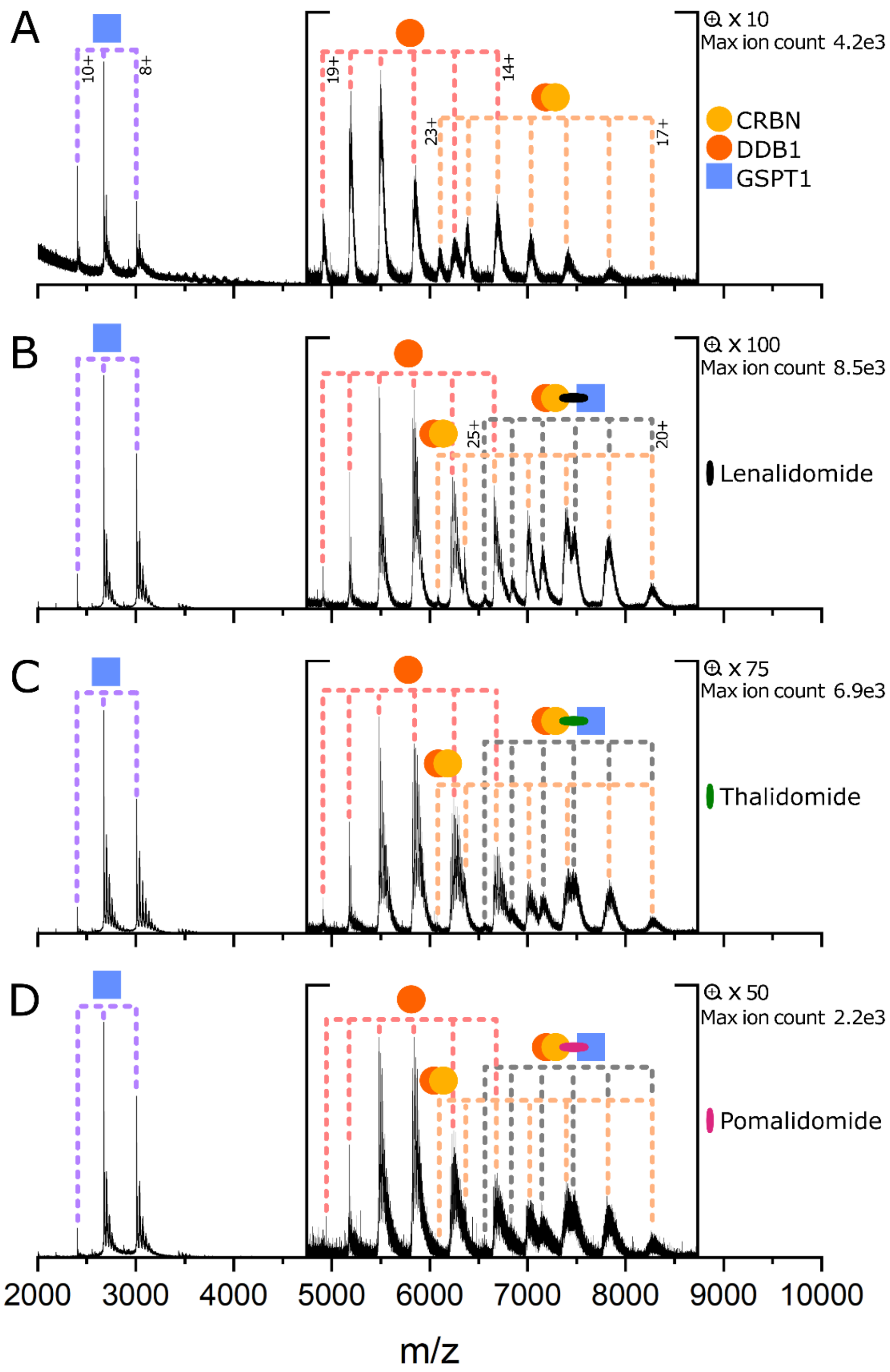


**Figure S1.** GSPT1 (5 µM) and CRBN:DDB1 (5 µM) in the absence of glue (A), presence of lenalidomide (100 µM) (B), thalidomide (100 µM) (C), and pomalidomide (100 µM) (D).

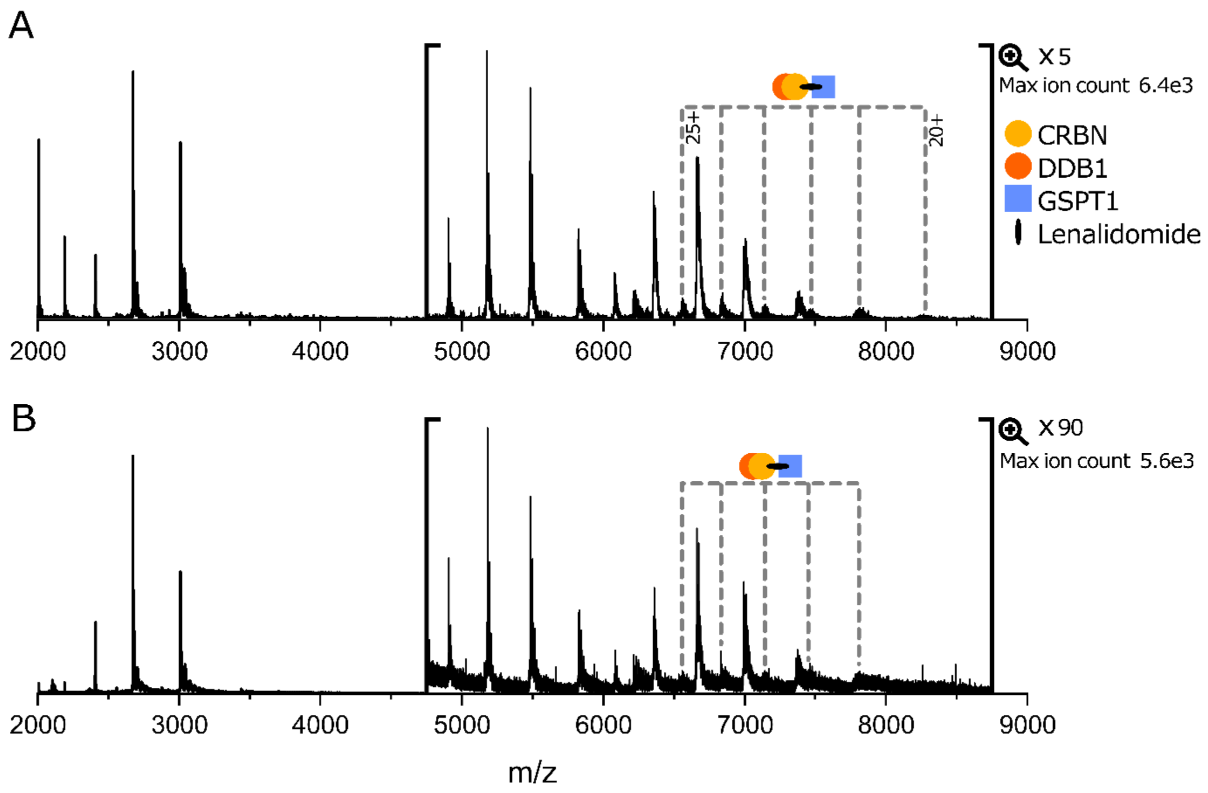


**Figure S2.** (A) GSPT1 + CRBN:DDB1 + lenalidomide (50 µM), (B) GSPT1 + CRBN:DDB1 + lenalidomide (5 µM). Protein concentrations are 5 µM and samples are in 100 mM AmAc containing 0.5% DMSO.

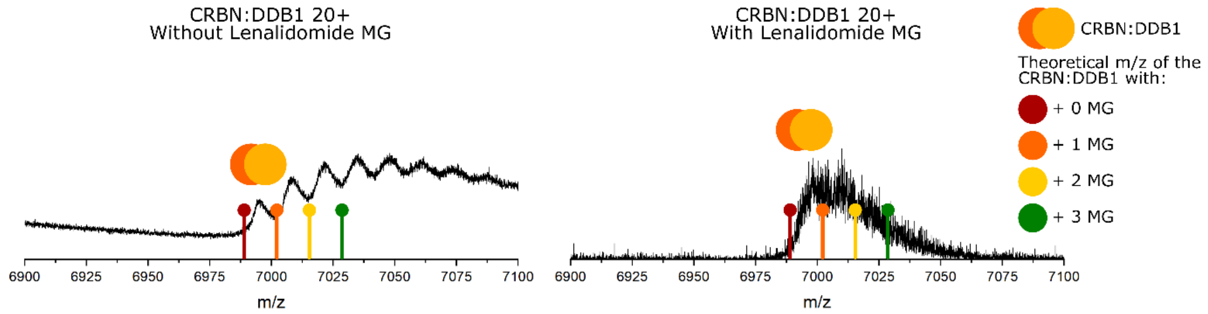


**Figure S3.** CRBN:DDB1 (5µM) in the absence and presence of lenalidomide (100 µM). The 20+ charge state peak has been focused on. The m/z at the leftmost of each peak is used to compare to the theoretical mass, as m/z values over the rest of the peak correspond to the species plus various adducts gained during desolvation. The m/z of both peaks aligns with the theoretical m/z of CRBN:DDB1 with no MG bound (6990 m/z). This shows no binary E3:MG species is observed upon addition of molecular glue. The adducts on the peak without Lenalidomide correspond to a molecular mass of ~280 Da, which is likely to be a detergent used in protein purification.

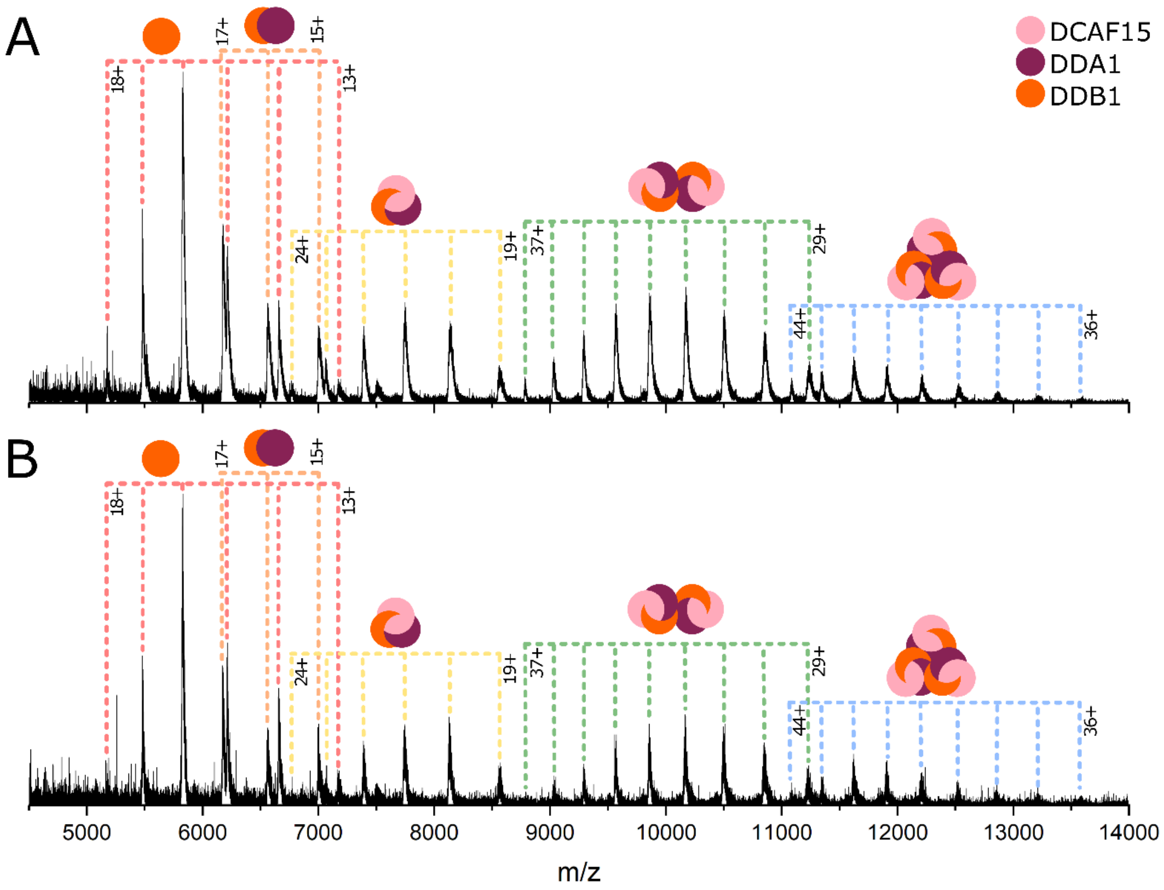


**Figure S4.** DCAF15 complex at 5 µM (A) and 2.5 µM (B). All species are present at the same relative intensity at both concentrations, with an overall reduction in intensity for the lower concentration, as expected.

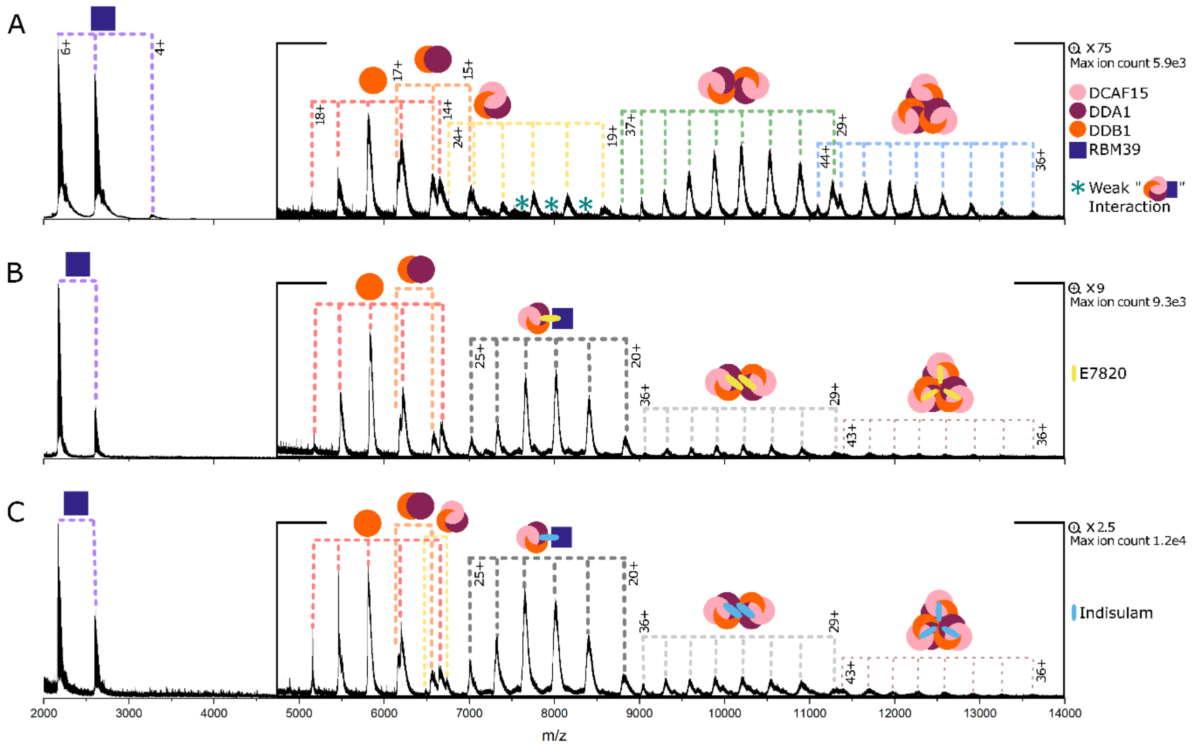


**Figure S5.** Mixture of RBM39 (5µM) and DCAF15 complex (5 µM) in the absence (A) and presence of E7820 (100 µM) (B) and indisulam (100 µM) (C).

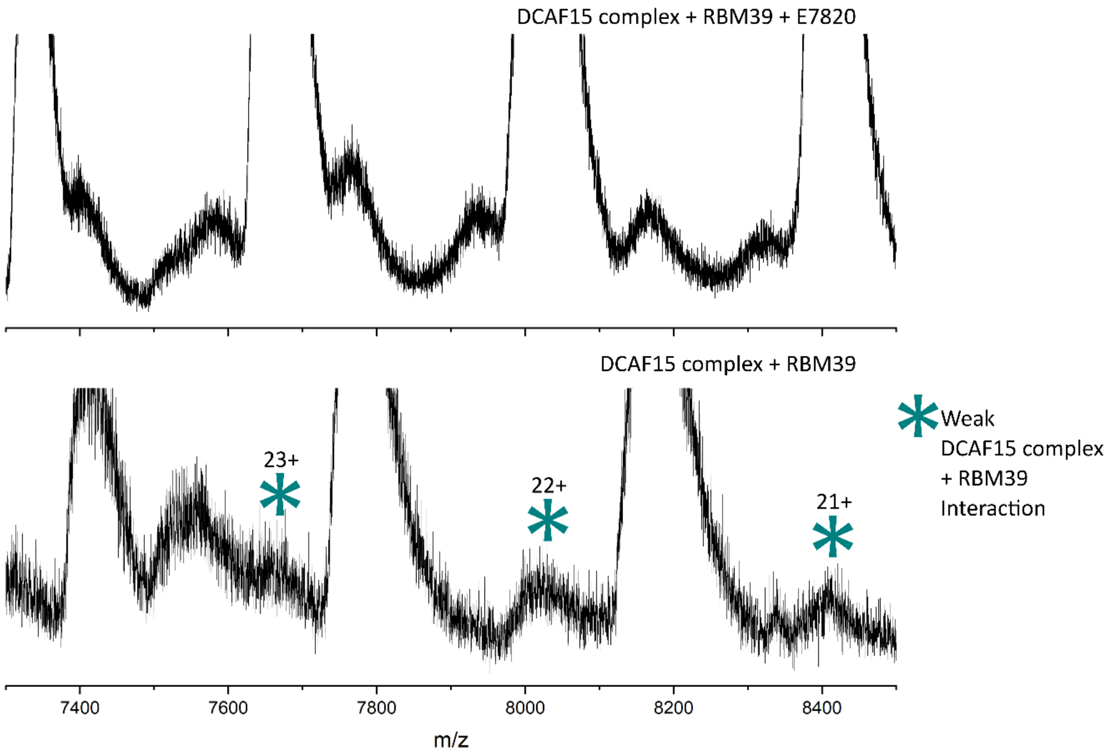


**Figure S6.** Mixture of RBM39 (5µM) and DCAF15 complex (5µM) in the presence (top) and absence of E7820 (100 µM) (bottom) showing low intensity peaks corresponding to the weak DCAF15 complex RBM39 interaction. These peaks cannot be observed in the spectrum containing molecular glue as the have a m/z value extremely close to that of the ternary complex, causing these small peaks to be masked by ternary complex peaks.

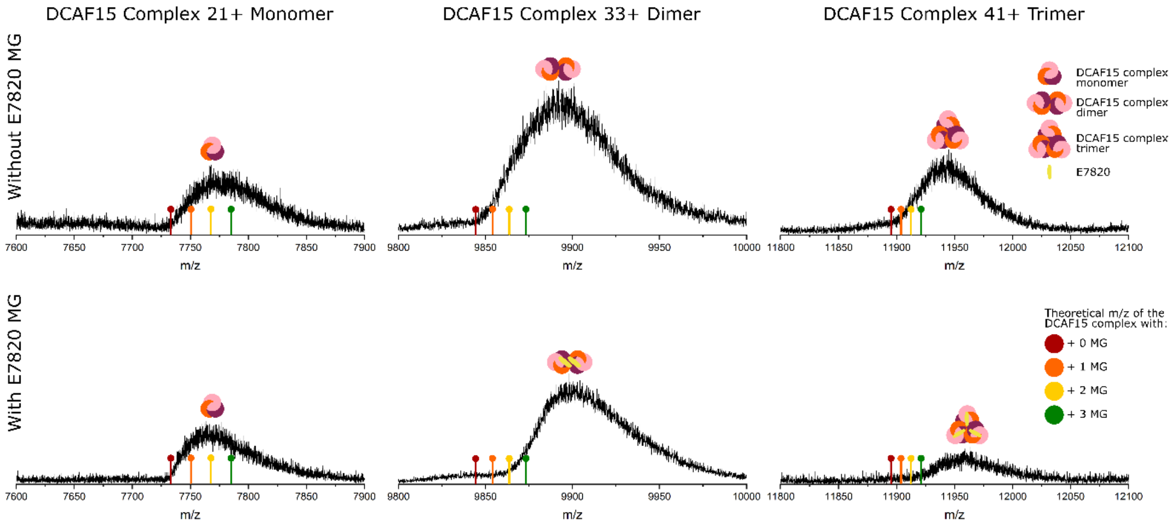


**Figure S7.** Selected peaks of RBM39 (5μM) and DCAF15:DDA1:DDB1 (5μM) in the absence (top) and presence (bottom) of E7820 (100 μM). Full spectra can be viewed in Figures 3A and C respectively in the main text. A mass shift can be observed for the dimer and trimer following E7820 addition, showing dimers and trimers of the DCAF15 complex do bind to the MG. The shift for the dimer is 20 m/z, which in charge state 33+ corresponds to a mass difference of 660 Da, approximately the molecular weight of two E7820 molecules (672 Da theoretical mass increase). The shift for the trimer is 23 m/z, which in charge state 44+ corresponds to a mass difference of 1012 Da, approximately the molecular weight of three E7820 molecules (theoretical mass shift 1008 Da).

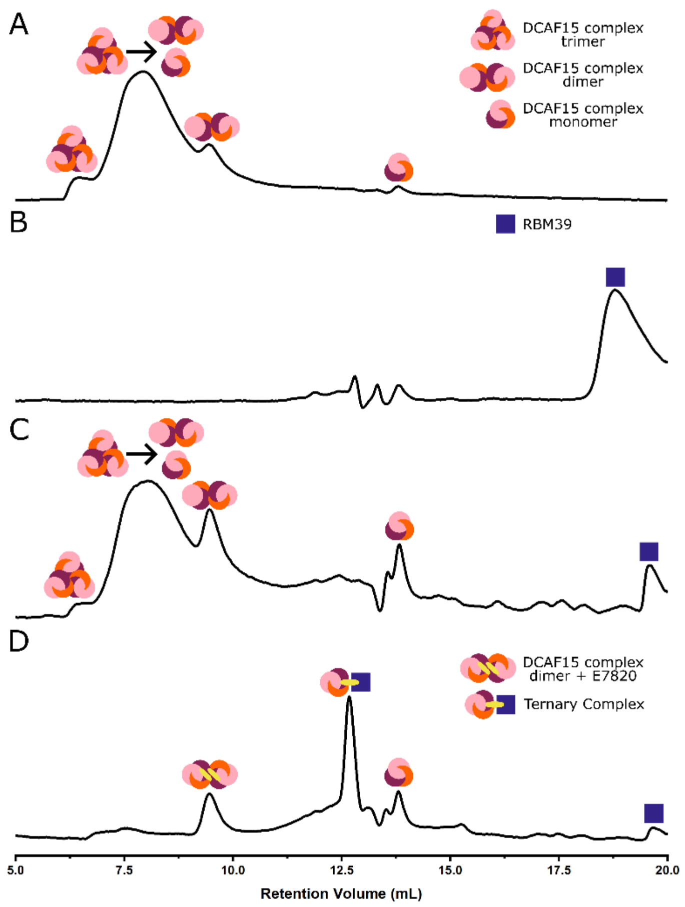


**Figure S8.** Size exclusion chromatography of (A) DCAF15 complex, (B) RBM39, (C) DCAF15 complex + RBM39, (D) DCAF15 complex + RBM39 + E7820.

(A)- DCAF15 complex alone. Peaks at retention volumes of 6.5 and 9 mL are attributed as trimeric and dimeric DCAF15 complex, respectively. The large, broad peak between 7 and 9 mL is attributed as being products of trimer dissociation (dimer and monomers) that has occurred during the time course of the experiment. This would also account for the raised baseline after the dimer (i.e., after 10 mL), as this would be the dissociated monomer. The monomer of the DCAF15 complex arrives at 13.5 mL.

(B)- The peak eluting at 18-19 mL is assigned as unbound RBM39.

(C)- DCAF15 complex + RBM39 in the absence of E7820. There is a similar distribution of the DCAF15 complexes existing as dimers and trimer.

(D)- DCAF15 complex + RBM39 + E7820. There are changes to the peak distribution that is in agreement with the findings from nMS. Most importantly, a new peak is observed at 12.5 mL which is assigned as the 1:1:1 DCAF15:E7820:RBM39 complex. There is also a large reduction in the intensity of the DCAF15 trimer peak, as well as the peak corresponding to the dissociation products of the trimer.

### Supplementary Tables

**Table S1.** Structures and molecular weights of all glues used in the study.

| Name | Structure | Molecular Weight (Da) |
|---|---|---|
| Lenalidomide | 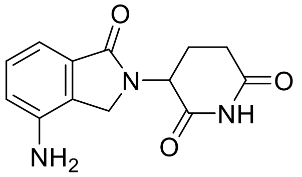 | 259.2 |
| Thalidomide | 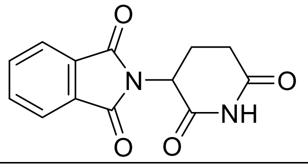 | 258.2 |
| Pomalidomide | 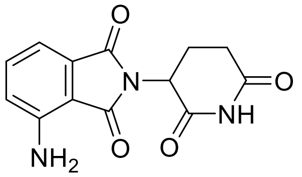 | 273.2 |
| Indisulam | 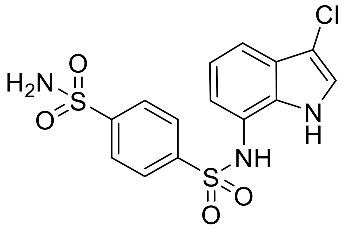 | 385.8 |
| E7820 | 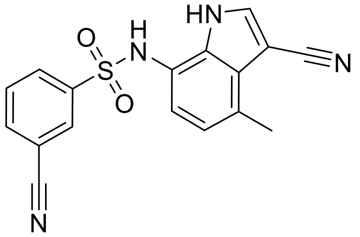 | 336.4 |

**Table S2.** Theoretical and measured molecular weights of proteins used in the study.

| Protein | Theoretical MW from sequence (Da) | Measured MW (Da) |
|---|---|---|
| CRBN_DDB1 | 139 772.0 | 139 775.5 ± 5.09 |
| GSPT1 | 23 832.8 + 226 (biotin) = 24 058.8 | 24 057.7 ± 0.20 |
| DCAF15 complex | 162 404.7 | 162 392.3 ± 18.00 |
| RBM39 | 12 758.4 + 226 (biotin) = 12 984.4 | 12 978.0 ± 0.01 |

Measured mass is the average from all labelled charge states, and the error is the standard deviation.
